# Rumen Sampling Methods Bias Bacterial Communities Observed

**DOI:** 10.1101/2021.09.22.461352

**Authors:** Jill V. Hagey, Maia Laabs, Elizabeth A. Maga, Edward J. DePeters

## Abstract

The rumen is a complex ecosystem that plays a critical role in our efforts to improve feed efficiency of cattle and reduce their environmental impacts. Sequencing of the 16S rRNA gene provides a powerful tool to survey shifts in the microbial community in response to feed additives and dietary changes. Oral stomach tubing a cow for a rumen sample is a rapid, cost-effective alternative to rumen cannulation for acquiring rumen samples. In this study, we determined how sampling method, as well as type of sample collected (liquid vs solid), bias the microbial populations observed. The abundance of major archaeal populations was not different at the family level in samples acquired via rumen cannula or stomach tube. Liquid samples were enriched for the order WCHB1-41 (phylum Kiritimatiellaeota) as well as the family *Prevotellaceae* and had significantly lower abundance of *Lachnospiraceae* compared with grab samples from the rumen cannula. Solid samples most closely resembled the grab samples; therefore, inclusion of particulate matter is important for an accurate representation of the rumen microbes. Stomach tube samples were the most variable and were most representative of the liquid phase. In comparison with a grab sample, stomach tube samples had significantly lower abundance of *Lachnospiraceae*, *Fibrobacter* and *Treponema.* Fecal samples did not reflect the community composition of the rumen, as fecal samples had significantly higher relative abundance of *Ruminococcaceae* and significantly lower relative abundance of *Lachnospiraceae* compared with samples from the rumen.

## Introduction

The ruminant stomach consists of four chambers the reticulum, rumen, omasum, and abomasum. The rumen, which is the largest of the four compartments, is a complex pregastric anaerobic fermentation chamber that harbors a diverse microbial community of bacteria, archaea, protozoa, and fungi [1]. These microbes exist symbiotically inside the ruminant host and are responsible for fermentation of dietary compounds. During the anaerobic fermentation of chemical constituents in the diet, volatile fatty acids (VFA), B-vitamins, and microbial cell proteins are produced, which serve as sources of nutrients and energy for the host that have a direct effect on physiological and production parameters [2]. As the rumen compartment does not secrete enzymes, ruminants are dependent on the enzymes produced by the various rumen microbes for digestion of feed. These microbial enzymes allow the ruminant to convert a wide variety of both plant- and animal-based feedstuffs into products that will contribute to the synthesis of meat and milk for human consumption. The bacterial population of the rumen comprises nearly 95% of the total microbial community and is diverse. There are many genera of bacteria that have been linked to feed efficiency, milk yield, and milk composition in dairy cattle [3,4].

Factors such as age [5], breed [6–8], health status, season [9] and diet of the animal all contribute to variation in the microbiota of the rumen. Dietary composition was reported to be the primary factor affecting the taxa present in the rumen microbiota as well as the richness of those taxa, with the ratio of forage-to-concentrate in the diet of utmost importance [10–12]. The rumen is home to a stable yet dynamic microbial ecosystem that has adapted to survive in an anaerobic environment with osmotic pressure, high buffering capacity and internal competition for substrate [13]. When dietary changes occur slowly, rumen conditions change causing microbial populations to shift in response to the new feed ingredients by favoring the growth of certain taxa over others, which subsequently affects the organic acid profiles produced [14]. However, when a dietary change occurs rapidly, for example changing from high forage diet (high cellulose and hemicelullose substrate) to a high concentrate diet (high starch and sugar), the shift in microbial community often causes simple indigestion in the cow, which can lead to ketosis. This occurs in dairy production when cows transition from a high forage diet fed prepartum to a lower forage, higher concentrate lactation diet in a matter of hours, which can contribute to indigestion. Management strategies at the farm level have evolved to minimize perturbations to the rumen microbial environment that reduce health and production performance when ingredients in the diet change due to cost or availability of a feed ingredient. Methods to quickly sample and diagnose microbial perturbations due to dietary transitions could improve the management strategies of these high-risk animals.

The composition of the rumen microbiota was first described by Hungate in 1966, and has been studied more extensively in recent years, in part due to the reduced costs associated with next generation sequencing techniques such as the pyrosequencing and Illumina platforms [15]. Much of the recent interest in the rumen microbiota has been generated by research related to climate change and the potential to reduce methane emissions from ruminant livestock as a greenhouse gas mitigation strategy. Next generation sequencing has thus far been a successful tool for characterizing the diversity of the microbial community within the rumen in greater detail through 16S rRNA gene amplicon profiling [16,17]. This technology is advantageous in that it allows the identification of a broader array of rumen microbial taxa, given that only a small fraction of the total species have been successfully cultured. However, the most appropriate method of obtaining a representative rumen sample is still widely debated [18]. It is well known that the bacterial populations between the solid and liquid portions of the rumen digesta differ in microbial composition, suggesting that the sampling method used will affect the characterization of the microbial community [19–24]. Thus, identifying sampling methods that accurately represent both the liquid and solid fractions of the rumen digesta are necessary.

Much of the existing research describing the rumen microbiome was performed on animals surgically fitted with rumen cannula, which offer the accuracy and convenience of sampling both liquid and solid rumen digesta directly from the rumen chamber. However, the surgical fistulation procedure is invasive, and the costs associated with the procedure as well as the ongoing animal care limit the number of animals that can feasibly be used in an experiment. Importantly, if microbial biomarkers of health or disease are identified for on-farm testing, retrieving rumen fluid through a cannula is not a practical approach on commercial dairy and livestock farms. Alternatively, many studies have used an oral stomach tube to collect rumen fluid without the need for a rumen fistula [8,18,25]. Oral stomach tubes are a cheaper, less invasive approach to rumen sampling that can be performed on as many cows as necessary, thus economically increasing the experimental sample size. In terms of bacterial community composition and diversity, rumen fluid extracted via the fistula was comparable to fluid extracted via the oral stomach tube [8,26]. Some of the disadvantages of the oral stomach tube include possible contamination by saliva (which affects the pH of the sample), inconsistent sampling region within the rumen, stress to the animal, skilled labor associated with use, and limited representation of particulate matter in samples, though the importance of these concerns to the microbial composition of the sample are widely debated among researchers [18,27].

The collection of fecal material from cattle is another non-invasive, simple, and inexpensive technique that is not as commonly regarded as a viable tool for collecting samples representative of the rumen microbiota. Although fecal sampling requires minimal equipment, is cost-effective, and can be performed easily on any animal, bacterial populations of the feces were found to not reflect the rumen digesta [28,29]. However, in these studies, the fecal microbiome was not compared with the liquid and solid fractions of the rumen digesta individually. If the feces reflect the microbial populations in the solid fraction, fecal samples might be useful in evaluating microbial taxa involved in fiber digestion. Conversely, if fecal samples represent the liquid fraction, lactate-producing microbes that contribute to ruminal acidosis could be diagnosed in a less invasive manner.

The aim of this study was to identify and compare the bacterial populations present in samples collected using three methods – an oral stomach tube, fecal samples, and grab sample through a rumen fistula. To the authors’ knowledge, no studies have considered this variety of sampling methods on a comparative basis using next generation sequencing. Our results will be useful in helping investigators design experiments that capture their microbial populations of interest.

## Materials and methods

### Animals

The experimental protocol and all procedures used in this study were approved by the UC Davis Institutional Animal Care and Use Committee. Four non-lactating Holstein (3) and Jersey (1) cows, each ruminally fistulated prior to the study, were used for the collection of samples. For the two-week duration of the study, cattle were housed individually with ad libitum access to water and offered the same maintenance total mixed ration (TMR) twice daily at approximately 08:00 and 16:00. Dietary composition of TMR was analyzed for protein, fiber, mineral, and energy content (Cumberland Valley Analytical Services, Hagerstown, MD; Table 1).

**TABLE 1.**
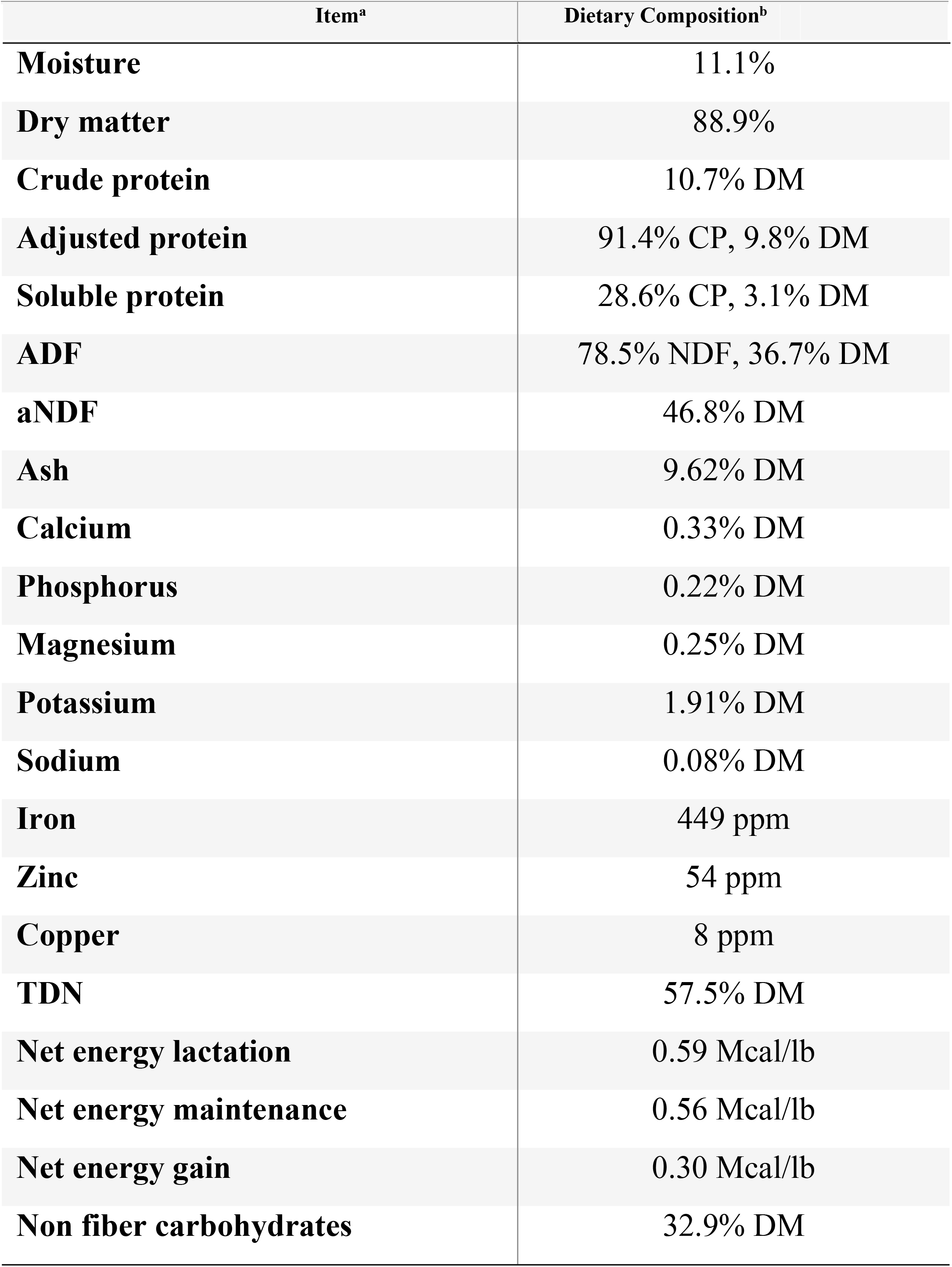
Dietary Composition of Total Mixed Ration. Chemical composition of the total mixed-ration (TMR) fed to the rumen-fistualted dry cows. Dietary analysis conducted by Cumberland Valley Analytical Services (Hagerstown, MD) completed 12/01/2016. Ingredient composition of TMR on an as is a basis was 50% wheat hay, 25% alfalfa hay, 21.4% almond hulls and 3.6% mineral supplement. ^a^Acid detergent fiber (ADF); Ash free Neutral Detergent Fiber (aNDF); Total Digestible Nutrients (TDN). ^b^Dry Matter (DM); Crude Protein (CP); Neutral Detergent Fiber (NDF).

### Sampling

Cows were given a one-week period for environmental adaption prior to sampling. This adaptation period was necessary to allow them to acclimate to an individual (rather than group) feeding approach and to reduce sorting of the feed. All cows were fed the maintenance TMR diet (Table 1) prior to and throughout the study. Sampling of fecal and ruminal contents occurred on days 7, 9, and 11 of the experiment, and took place approximately 4 hours after morning feeding. Fecal samples were collected from the rectum with sterile polyethylene gloves and stored in plastic bags. Grab samples (containing both liquid and particulate matter) from the fistula were collected from the medioventral region of the rumen and stored in plastic bags. Rumen liquid was collected from the fistula using a PVC pipe, Tygon® tubing, and a large syringe, and stored in 240 ml sterile plastic vials. The Tygon® tubing was thoroughly rinsed and bleached between cows to avoid cross-contamination of samples. For liquid strained samples, about 250 ml of liquid sample was squeezed through 4 layers of cheesecloth to remove large particles, as is common done [19,30,31]. For solid samples, similar squeezing through a cheese cloth was applied to remove liquid from the solid digesta content, before being stored in plastic bags. On days 9 and 11, the first aliquot of rumen liquid, containing both liquid and solid particulates, was additionally collected as a liquid unstrained sample in 240 ml sterile plastic vials. Lastly, enough rumen liquid to fill a 240 mL sterile plastic vials, which was collected via an oral stomach tube using an oral speculum, Tygon® tubing (1.5cm O.D. and 0.9cm I.D.) and a vacuum pump. A fresh tube was used for each cow to avoid cross-contamination of samples. The pH of each of the liquid-containing samples was measured with a portable pH meter (Milwaukee Instruments, Rocky Mount, NC). All samples were held on ice during transport and stored in triplicate 60 ml vials at −20°C for DNA extraction and dry matter analysis.

### DNA extraction and PCR amplification

DNA extraction was performed using a ZR Fecal DNA MiniPrep™ kit (Zymo Research Corp., Irvine, CA), with slight modifications to the manufacturer’s instructions. Samples were thawed at room temperature, and 200 mg of each sample were used for DNA extraction, which included a bead bashing step to facilitate the mechanical lysis of microbial cell walls. As the last step in the procedure, DNA was eluted from the column with elution buffer, and the resulting DNA was evaluated for concentration and purity on a NanoDrop 2000 spectrophotometer (Thermo Scientific, Waltham, MA, USA) and stored at −20°C. The V4 region of the bacterial 16S rRNA gene was amplified from each sample using forward primer F515 containing a unique 8 bp barcode (N) and linker region (**GT**) (5’-NNNNNNNN**GT**GTGCCAGCMGCCGCGGTAA-3’) and the reverse primer R806 (5’-GGACTACHVGGGTWTCTAAT-3’). The amplification was carried out in triplicate using GoTaq® Green Master Mix (Promega, Madison, WI) as previously described [32]. In brief, PCR conditions were set at initial denaturation for 94°C for 3 min; followed by 35 cycles of 94°C for 45 seconds, 50°C for 1 min, 72°C for 90 seconds with final extension step at 72°C for 10 min. Triplicates were combined in equal concentrations and amplicons were evaluated for off target bands by gel electrophoresis, pooled and then purified using a QIAquick PCR Purification Kit (QIAGEN, Hilden, Germany). A 50µl aliquot of the final pooled PCR product was sequenced at the UC Davis Genome Center DNA Technologies Core via the Illumina MiSeq PE250 platform (Illumina, CA).

### Amplicon library processing

Raw paired end reads were screened to remove phiX, human and host contamination using Kneaddata v0.6.1 by aligning reads to the phiX174 (NCBI ACC: NC_001422.1), bovine (ARS-UCD1.2) and human (GRCh38) reference genomes [33]. Reads were demultiplexed followed by trimming of primers and barcodes with Cutadapt v1.18 [34]. Ends of reads were trimmed for quality, any read smaller than 150bp was discarded and a max expected error of 2 was used as a quality filter using the filterAndTrim function from DADA2 v1.8.0 [35]. Sequences were merged, denoised, chimeras were removed and exact amplicon sequence variants (ASVs) were identified using DADA2. Taxonomy was assigned using the RDP native Bayesian classifier algorithm in the DADA2 assignTaxonomy function with Silva reference database v.132 training set. A phylogenetic tree of unique ASVs was made using FastTree with default options in QIIME v.1.9.1 [36]. The ASV table, sequences and tree produced by DADA2 were imported into the R package Phyloseq v.1.24.2 for further analysis [37].

### Microbial community analyses and statistics

First, unsupervised exploratory analysis was conducted with double principal coordinates analysis (DPCoA), which was calculated and graphed with the phyloseq R package [37,38]. Both modeling and hypothesis testing of differentially abundant ASVs between sample types was determined using the Corncob R package [39]. All genera-level and ASV-level relative abundances were modeled using a beta-binomial regression with a logit-link for mean and dispersion as described by Martin et al [39]. Differential abundance was modeled as a linear function of sample type, cow and day. Significant differentially abundant ASVs were determined with the parametric Wald test with bootstrapping (n=1000) as described by Martin et al [39]. Within the Corncob algorithm the Benjamini-Hochberg (BH) adjustment for multiple comparisons was used to calculate adjusted *p* values. An adjusted *p* value ≤ 0 .05 was considered significant. This model has the benefit of accommodating the absence of a taxon in samples without zero-inflation or pseudocounts, accounts for differences in library sizes, give valid inference even with small samples [39]. Richness of sample types was estimated with the R package breakaway and evenness was calculated using the R package DivNet, which accounts for the structure of microbial communities [40,41]. Hypothesis testing of alpha diversity (richness and evenness) metrics was done using the betta() function using sample type, cow and day as fixed effects in the breakaway R package [42]. Beta diversity was calculated by using unweighted UniFrac distances and graphed by PCA clustering in the Phyloseq R package [37]. The number of clusters in the data was determined with the gap statistic using the gapstat_ord() function in Phyloseq [43].

### Data availability

Scripts for sequence processing and analysis, interactive graphs, R objects as well as an Rmarkdown file to reproduce figures in this paper can be found at https://doi.org/10.5281/zenodo.4026849. Raw sequencing files are available through the Sequence Read Archive under the study accession number PRJNA692782.

## Results

### Sequence processing of rumen and fecal samples

After filtering with Kneaddata and demultiplexing the single run of MiSeq yielded 747,961 250bp raw paired-end reads that entered the DADA2 pipeline. After the quality trimming, initial filtering, and chimera removal, the library size ranged from 2,189 to 24,624 reads, with a median library size of 7,197 and an average size of 8,110 reads. The median read length of quality filtered merged reads was 257bp. A total of 5,607 AVSs were identified, of which 94 weren’t assigned to a phylum and thus were removed for analysis along with 12 ASVs assigned to chloroplasts and mitochondria. The 94 unassigned taxa were found in all sample types with solid samples having the most reads of unknown taxa. This suggests there are is still a diverse group of microbes attached to solid particles that have yet to be identified. The final feature table had 5,485 ASVs across 68 samples.

### Community composition of all sample types

The 5,485 ASVs were assigned to 21 phyla, 78 orders, 117 families, and 293 genera. Here we define major phyla as those with a mean relative abundance in at least one sample type of greater than 3%. Major phyla were Firmicutes, Bacteroidetes, Kiritimatiellaeota, Proteobacteria, Euryarchaeota and Spirochaetes (Fig 1A). Of these major phyla, Firmicutes was significantly lower in relative abundance (*P* ≤ 0.0001; Fig 1C) while Bacteroidetes and Proteobacteria had significantly higher relative abundance in feces and liquid samples as compared with grab samples (*P* ≤ 0.002; Fig 1C). In addition, Kiritimatiellaeota was significantly higher in relative abundance in stomach tube and liquid samples compared to grab samples (*P* ≤ 0.001; Fig 1C). Spirochaetes was significantly lower in relative abundance in feces, stomach tube and solid samples compare with grab samples (*P* ≤ 0.003; Fig 1C). While Euryarchaeota had significantly lower relative abundance in feces, it had significantly higher relative abundance in stomach tube samples compared with grab samples (*P* = 3.24×10^−8^). Minor phyla were those with less than 3% relative abundance in all samples (Fig 1B). An interactive version of Figure 1B with mean and standard deviations for each phyla is available at https://doi.org/10.5281/zenodo.4026849 as interactive Fig 1 – minor phyla. The phylum Gemmatimonadetes was only found in stomach tube samples and Deferribacteres was only found in fecal samples (Figure 1B). For the minor phyla in feces Tenericutes, Patescibacteria, Actinobacteria, Fibrobacteres, Chloroflexi and Synergistetes were significantly lower in relative abundance and Verrucomicrobia, Epsilonbacteraeota, Cyanobacteria, Planctomycetes and Lentisphaerae were significantly higher in relative abundance compared with grab samples (*P* ≤ 0.001; Figure 1C). Samples acquired with the oral stomach tube had significantly lower relative abundance of Patescibacteria and Fibrobacteres and significantly higher relative abundance of Verrucomicrobia, Epsilonbacteraeota, and Fusobacteria compared with grab samples. Only 1.68% of ASVs were assigned a species, but 67.3% were able to be assigned to a genus.

**Fig 1.**
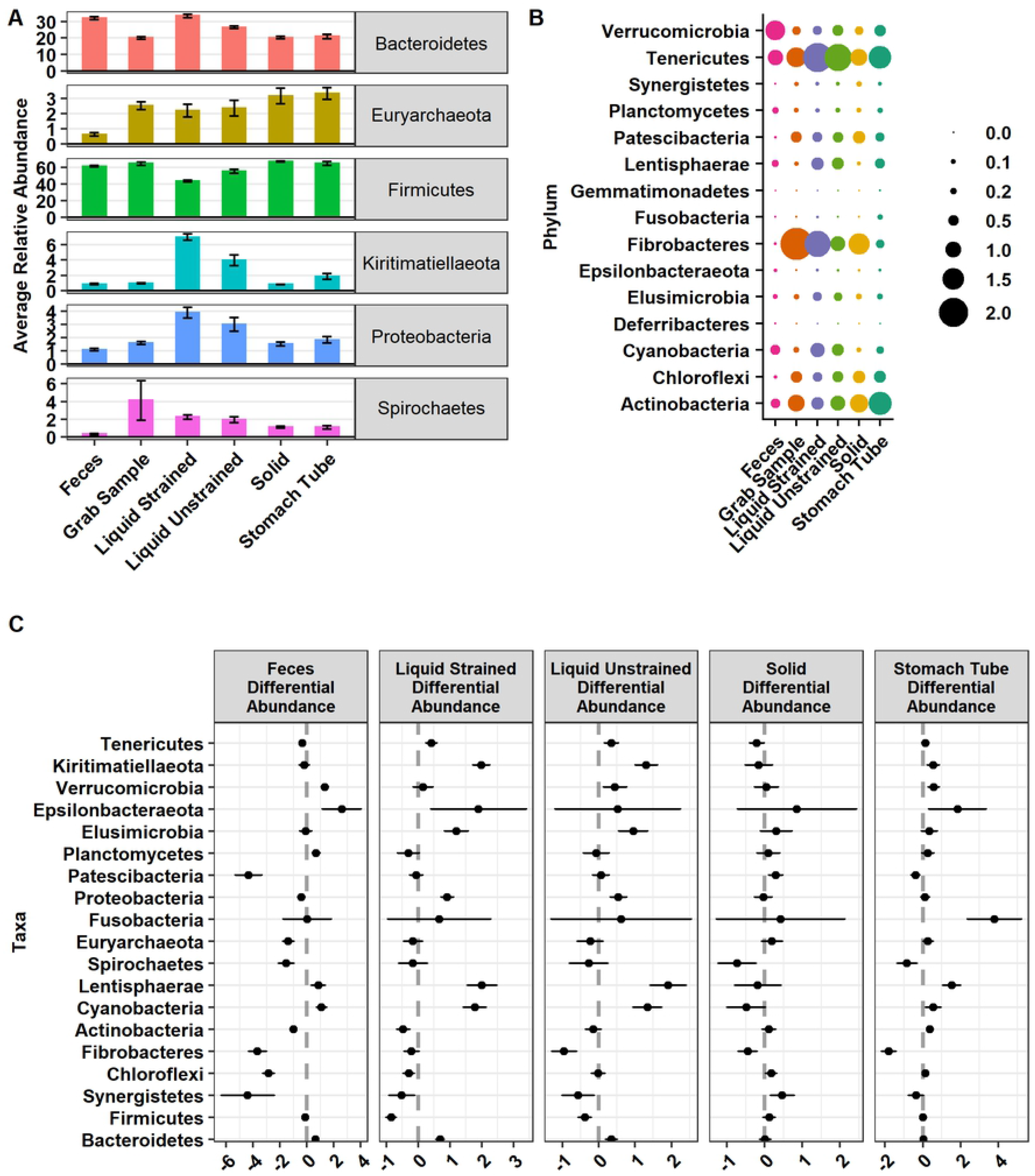
Relative abundance of (A) major and (B) minor phyla and (C) their differential abundances. (A) Relative abundance of major phyla defined as those phyla found at greater than 3% relative abundance and graphed as relative abundance ± SE. (B) Minor phyla defined as those found below 3% relative abundance present in sample types. (C) Phyla that are significantly differentially abundant compared with grab samples. Graphed as coefficients with a 95% confidence interval from the corncob model. Families with negative coefficients for a sample type are expected to have a lower relative abundance when compared to the grab samples while positive coefficients suggest a higher relative abundance in that sample type compared to grab samples.

Liquid unstrained and fecal samples were the least variable samples as they shared 510 and 441 ASVs, respectively, with samples of their own type. On the other hand, stomach tube and liquid strained samples were the most variable as these sample types only shared 225 and 307 ASVs, respectively, with samples of their own type. Moderately variable sample types were grab and solid samples, which shared 319 and 405 ASVs, respectively, with samples of their own type.

### Diversity

The evenness of fecal samples was significantly lower than all rumen sample types (*P* ≤ 0.001; Fig 2A). Fecal, stomach tube, and liquid strained samples had significantly lower evenness than grab samples (*P* ≤ 0.001; Fig 2A). Solid and liquid unstrained samples did not have significantly different evenness compared with grab samples (*P* ≥ 0.05; Fig 2A). Both the individual cow sampled and day of sampling significantly affected the evenness of a sample (*P* ≤ 0.05; Fig 2A).

**Fig 2.**
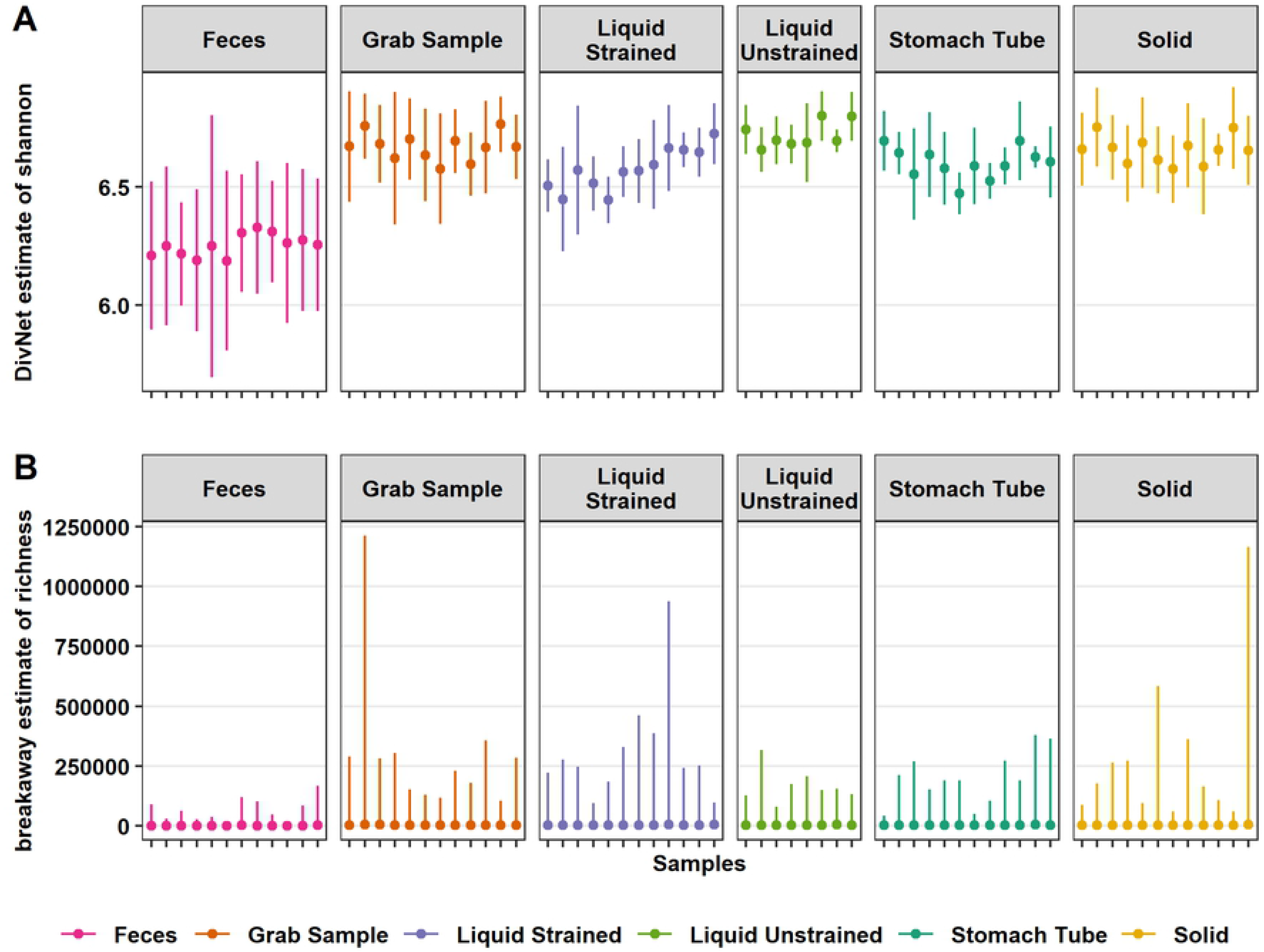
Differences in estimated alpha diversity among sample types. (A) DivNet estimate of Shannon diversity plotted as mean with a 95% confidence intervals and (B) mean breakaway estimate of species richness with 95% confidence intervals. Both the richness and evenness of fecal samples were significantly lower than all other rumen sample types (P ≤ 0.001). Stomach tube and Liquid strained samples had significantly lower evenness than grab samples (P ≤ 0.001). Solid and stomach tube samples were estimated to have significantly fewer species than grab samples (P = 0.02 and P ≤ 0.001, respectively).

The richness of samples from the rumen were estimated to be significantly higher than that of fecal samples (*P* ≤ 0.001; Fig 2B). Fecal samples were estimated to have a mean of 2,021 species, which was significantly lower than the grab samples estimated mean of 4,119 species (*P* ≤ 0.001; Fig 2B). Liquid strained and unstrained samples did not have a significantly different mean number of estimated species compared with grab samples (*P* ≥ 0.05; Fig 2B). However, solid and stomach tube samples were estimated to contain a significantly lower number of species compared with grab samples, 286 and 506, respectively (*P* = 0.02, *P* ≤ 0.001; Fig 2B). Neither the day sampled nor individual cow had a significant effect on the number of species in a sample (*P* ≥ 0.05; Fig 2B).

Weighted UniFrac distances were calculated to determine beta diversity. Calculations of eigenvalues showed that 86.8% of the variance between samples was contained in the first two principle components, thus a two-dimensional visualization was deemed appropriate (Fig 3). Two distinct groups were present with fecal samples clustering away from all rumen sample types (Fig 3). Grab and solid samples exhibited low variability and overlapped each other, forming one group. Liquid samples were further down the second axis, which might indicate that there were distinct phylogenetic differences between these samples and grab samples. Stomach tube samples were the most variable with some of these samples found within the grab and solid sample cluster, while other stomach tube samples were more closely associated with liquid samples. The gap statistic of the weighted UniFrac strongly suggested there were at least 3-5 clusters in the data. As there are six sample types in the dataset, this suggests that grab and solid samples are likely one cluster as these samples overlap the most (Fig 3). The unweighted UniFrac showed a similar pattern, with less variation explained in axis one and two, 45.8% and 6.9%, respectively (data not shown).

**Fig 3.**
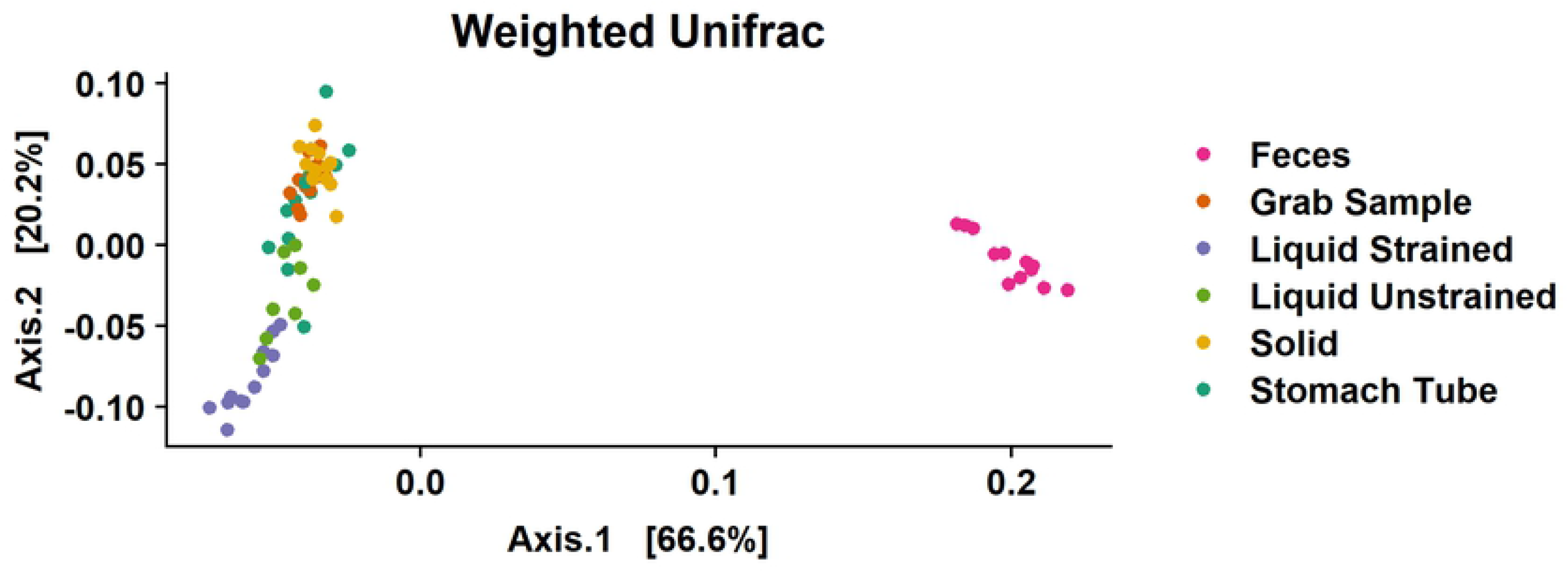
Beta diversity as weighted UniFrac distances between samples. To faithfully reflect the variance in the coordinates, the height-to-width ratio was based on the ratio between the corresponding eigenvalues.

### Overall differences between sample types

As an exploratory first step, DPCoA was performed (Fig 4A). An interactive version of this graph with taxon identification is available at https://doi.org/10.5281/zenodo.4026849 as interactive figure 2 – DPCoA. Additionally, since Firmicutes and Bacteroidetes dominated a majority of the graph, a version without these phyla was created at and is available at https://doi.org/10.5281/zenodo.4026849 as interactive Fig 3 – DPCoA_NoFrimBact with the aim to allow a better visualization of minor phyla. This phylogenetic ordination method provides a biplot representation of both samples and taxonomic categories. The DPCoA was used to identify the underlying structure of these data and identify taxa that could be contributing to differences between sample types that will be specifically examined with differential abundance testing.

**Fig 4.**
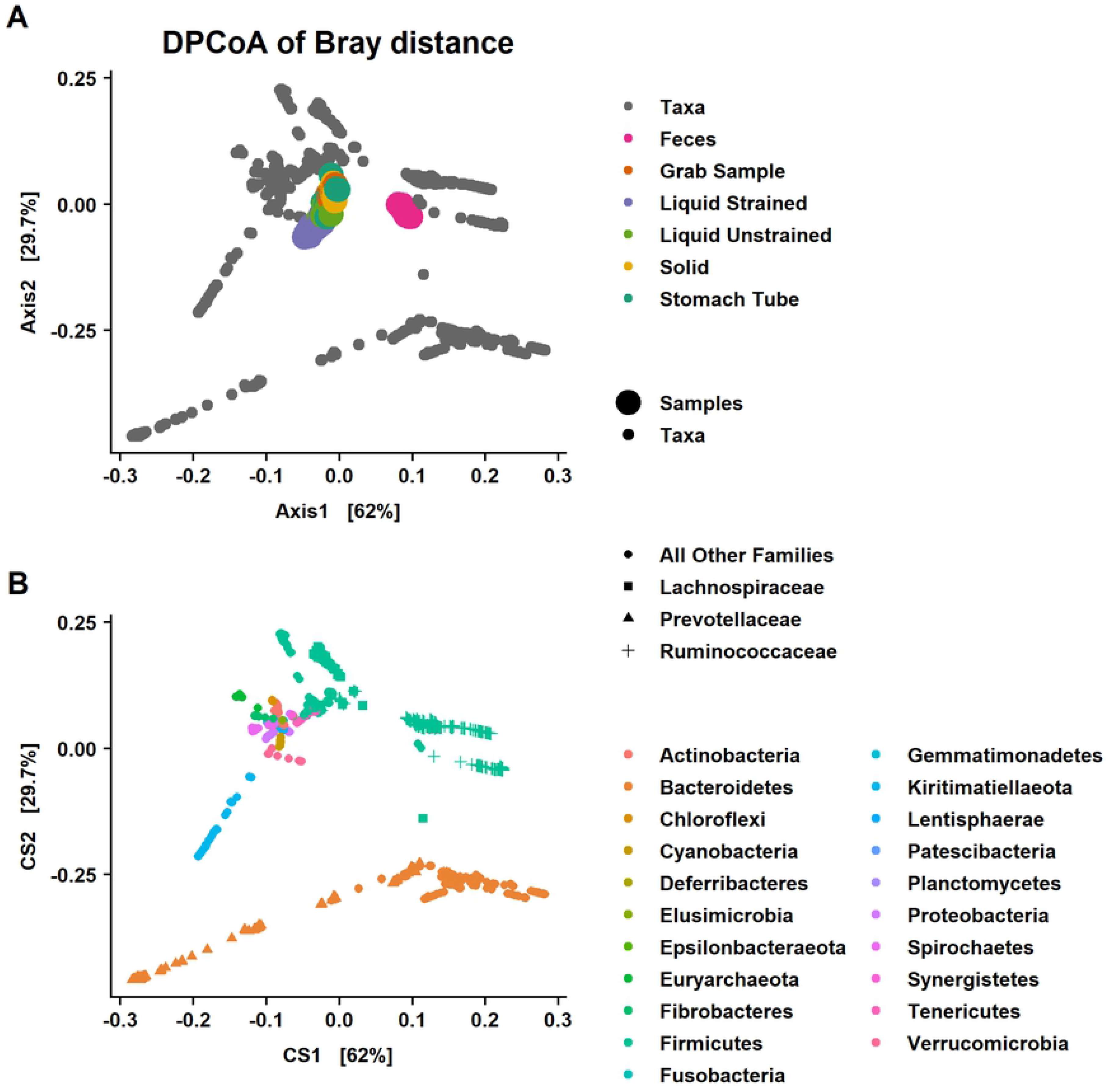
Double principal coordiant analysis (DPCoA) of the Bray-Curtis distances among samples. DPCoA is a phylogenetic ordination method and that provides a biplot representation of both (A) samples and (B) taxonomic categories. Note that while the biplots are a square shape to fit the page the CS1 explains roughly twice the variation of CS2 similar to what is seen in Fig 1. The 1^st^ axis discrimates fecal from rumen samples while the 2^nd^ axis separtes liquid strained samples from other rumen sample types. Samples that have larger scores on CS1 have a subset of taxa from Bacteroidetes and Firmicutes that is different than rumen samples. Liquid strained samples have lower values on CS2 suggesting they are distinguished from other rumen sample types by taxa in the phylum Kiritimatiellaeota and *Prevotellaceae*. Fecal samples are predicted to have lower abundance of *Lachnospiraceae* and greater abundance of *Ruminococcaceae*.

Fecal samples clustered away from samples that were collected from the rumen, which was primarily driven by differences in the relative abundance of a subset of Firmicutes in the families *Ruminococcaceae*, *Lachnospiraceae* and *Christensenellaceae* and a subset of families in the phylum Bacteroidetes, mainly *Rikenellaceae* and *Prevotellaceae* on the 1^st^ axis (Fig 4B). Additionally, fecal samples separate from samples from the rumen based on having more taxa from the family *Akkermansiaceae* and phylum Tenericutes and fewer from the families *Fibrobacteraceae*, and *Spirochaetaceae* (Fig 4B and S1 Fig). Liquid samples were found lower on the 2^nd^ axis of the DPCoA, indicating these samples had more taxa from the phylum Kiritimatiellaeota and a subset of Bacteroidetes most of them in the family *Prevotellaceae* (Fig 4B). Also, the separation of liquid strained samples away from other rumen samples was due to fewer taxa from the phylum Euryarchaeota and the family *Eggerthellaceae* that is within the phylum Actinobacteria (S1 Fig).

To test the significance of these differences, differential abundance testing was performed with Corncob. All sample types were compared to grab samples as a base line, because it is considered the gold standard for surveying microbial communities in the rumen. The relative abundance of *Prevotellaceae* was significantly lower in feces and was significantly higher in liquid samples compared with grab samples (*P* ≤ 0.0004; Fig 5A and 6). Stomach tube (*P* = 0.06) and solid (*P* = 0.77) samples were not significantly different in the relative abundance of *Prevotellaceae* compared with grab samples. The relative abundance of *Prevotellaceae* was highest in liquid strained samples compared with other sample types (Fig 5A). In comparison to grab samples, the relative abundance of *Ruminococcaceae* was significantly higher in feces (*P* ≤ 0.001; Fig 5B) and solid (*P* = 0.003; Fig 5B) samples while liquid strained samples had significantly lower relative abundance (*P* ≤ 0.001; Fig 5B). Neither stomach tube nor liquid unstrained samples had significantly different relative abundance of *Ruminococcaceae* compared with grab samples. Fecal samples were lower in relative abundance of *Lachnospiraceae* compared with all other samples (*P* ≤ 6.96×10^−15^), while relative abundance was higher for grab samples compared with all other sample types (*P* ≤ 0.03; Fig 5C). Neither day of sampling nor individual animal significantly affected the relative abundance of *Prevotellaceae* (*P* ≥ 0.05). In contrast, the relative abundance of *Ruminococcaceae* and *Lachnospiraceae* was significantly affected by individual animal (*P* ≤ 0.03), but not day of sampling.

**Fig 5.**
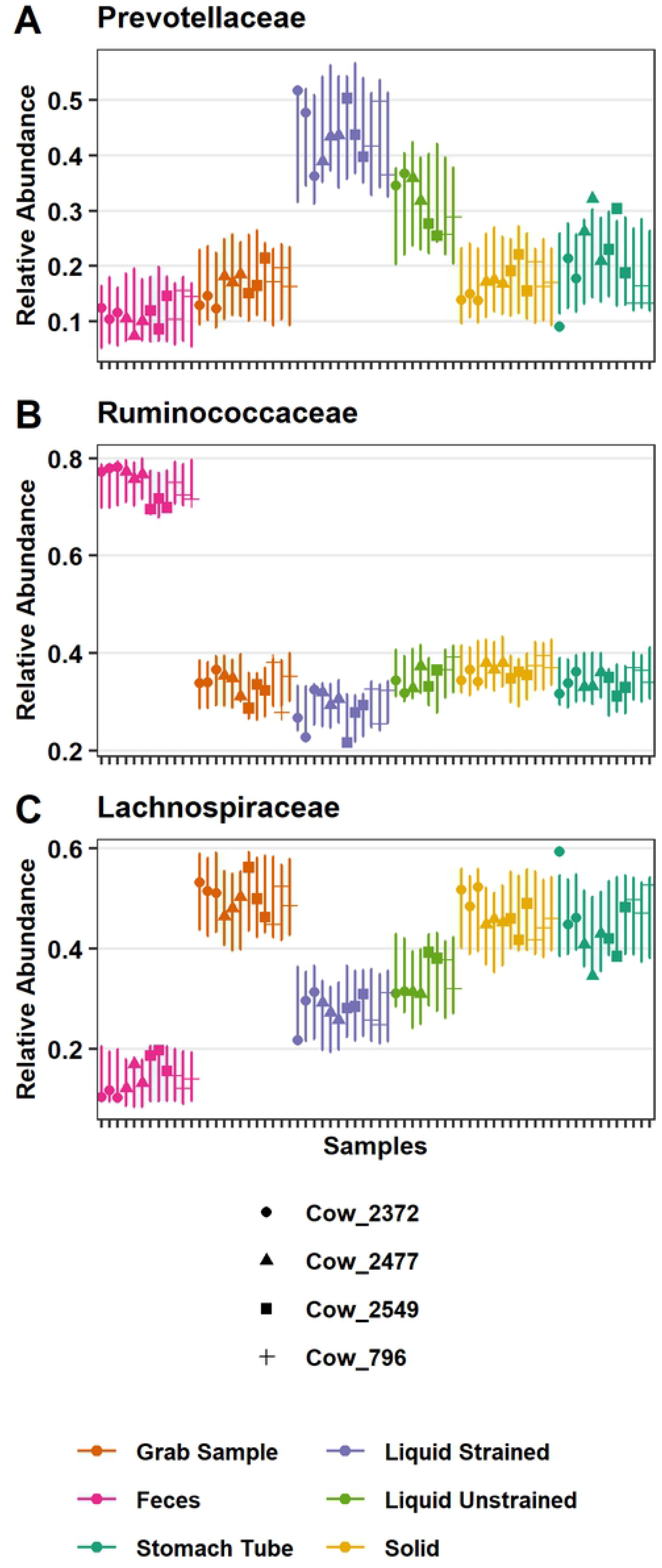
Significant differences in the relative abundance of specific bacterial families. Relative abundance of (A) *Prevotellaceae* (B) *Ruminococcaceae* and (C) *Lachnospiraceae* as modeled by corncob. Points are the estimated relative abundance and bars are a 95% prediction interval for each cow on different days of sampling.

**Fig 6.**
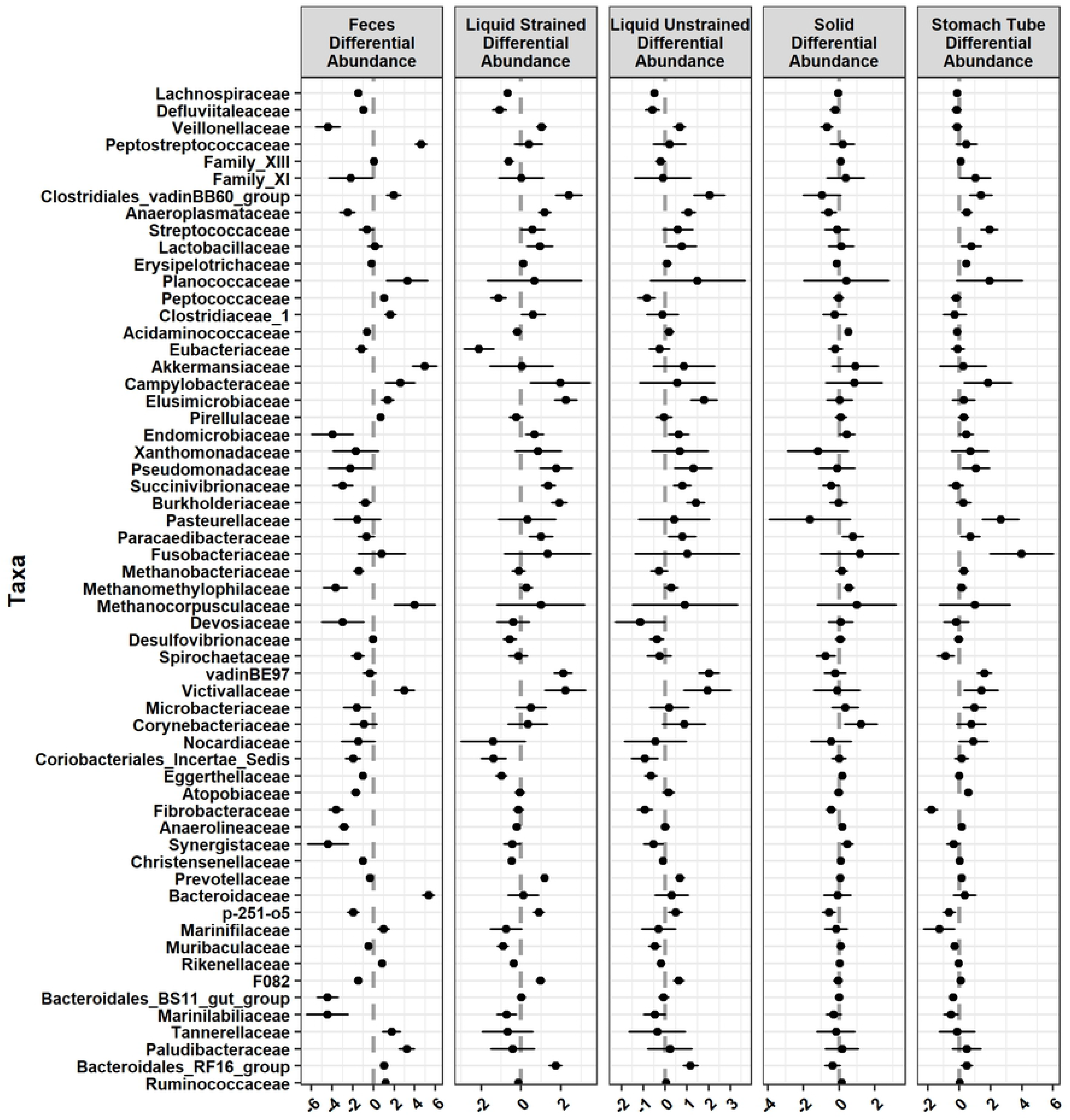
Families that were significantly differentially abundant across sample type compared with grab samples. Graphed as coefficients with a 95% confidence interval calculated from the corncob model. Families with negative coefficients for a sample type are expected to have a lower relative abundance when compared to the grab samples while positive coefficients suggest a higher relative abundance in that sample type compared to grab samples.

### Specific community differences between grab and fecal samples

To further distinguish what taxa were contributing to the separation of fecal samples from rumen samples on the DPCoA, we identified taxa that were found in one sample type and not the other. Within the phyla Firmicutes and Bacteroidetes, families *Barnesiellaceae*, *Chitinophagaceae*, *p-2534-18B5_gut_group*, *GZKB124*, and *Hymenobacteraceae* were found in fecal samples, but were not found in grab samples. Conversely, *Leuconostocaceae*, *Carnobacteriaceae*, *Aerococcaceae*, *Syntrophomoadaceae*, *Bacteroidetes_DB2-2*, *PeH15*, *M2PB4-65_termite_group*, *COB_P4-1_termite_group*, *Spirosomaceae*, and *Porphyromonadaceae* were found in grab samples, but were not found in fecal samples.

Next, we identified ASVs, genera and families that were differentially abundant between sample types. There were 657 significant differentially abundant ASVs in fecal samples compared with grab samples, as well as 114 differentially abundant genera (*P* ≤ 0.05; S2 Fig). At the genera level, 131 ASVs were unable to be fit to the Corncob model for differential abundance testing. Primarily, this was due to either limited or lack of reads in one of the sample types. Of these genera that didn’t fit the model, *Acetatifactor*, *Shuttleworthia*, *Succinivibrio, Veillonellaceae UCG-001*, and *Lachnospiraceae UCG-006* were found in all grab samples with greater than or equal to 50 reads across all samples, but absent in fecal samples. Similarly, there were 11 genera found in all fecal samples with 50 or more reads, but these were not found in any grab samples including *Coprococcus 3*, *Cellulosilytium*, *Clostridioides*, *Paeniclostridium*, *Parasutterella*, *Aeriscardovia*, *Odoribacter*, *Harryflintia*, *Negativibacillus, Pygmaiobacter*, and *Ruminococcaceae UCG-011*.

The most common families with differentially abundant ASVs were *Lachnospiraceae*, *Ruminococcaceae*, *Christensenellaceae*, *Family XIII, Rikenellaceae*, and *Prevotellaceae*. These families are in the phyla Firmicutes and Bacteroidetes, which had the most significant differentially abundant ASVs. However, as a percent of total ASVs these phyla only had 4.9% and 16.3% significant differentially abundant ASVs, respectively. In contrast, 25.6% of the ASVs assigned to Chloroflexi and 29.5% of ASVs assigned to Euryarchaeota were significantly different between grab and fecal samples. The significant ASVs in Chloroflexi were all assigned to the genus *Flexilinea*. In addition to the significantly lower abundance of some Chloroflexi ASVs in fecal samples compared with grab samples, another 51.3% of the ASVs in the phyla were not found in any fecal samples (S2 Fig). In the phylum Euryarchaeota, feces had significantly lower abundance of *Methanobrevibacter*, *Methanosphaera*, and were almost devoid of *Methanomethylophilaceae*.

There were 30 families that had significantly lower relative abundance while 18 families had higher relative abundance between fecal and grab samples (Fig 6). Families that had the strongest positive relationship with fecal samples were *Peptostreptococcaceae* (*P* = 1.76×10^−7^; Fig 7A), *Akkermansiaceae* (*P* = 6.95×10^−5^; Fig 7B), and *Bacteroidaceae* (*P* = 7.87×10^−12^; Fig 7C), which were significantly higher in relative abundance compared with grab samples. Conversely, the families with largest negative relationship between fecal and grab samples that had significantly lower relative abundance were *Veillonellaceae* (*P* = 1.66×10^−11^; Fig 7D) and Bacteroidales_BS11_gut_group (*P* = 6.20×10^−11^; Fig 7E). Additionally, fecal samples separated from rumen samples on the DPCoA (S1 Fig) due in part to differences in the families *Spirochaetaceae* and *Fibrobacteraceae* both of which had significantly lower relative abundance than grab samples (*P* = 7.88×10^−9^; *P* = 4.73×10^−8^, respectively Fig 7F).

**Fig 7.**
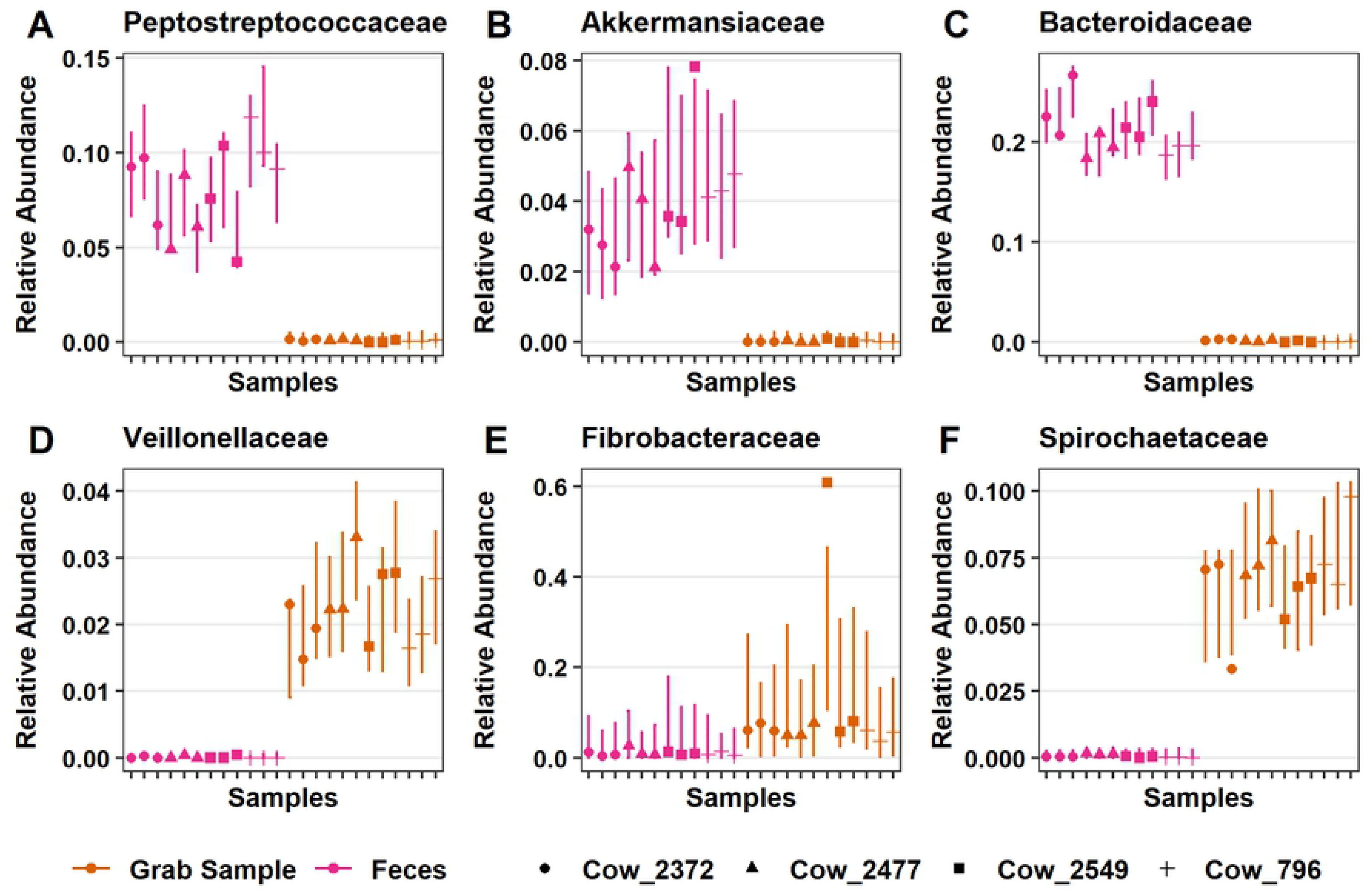
Significant differences in the relative abundance of specific bacterial families between fecal and grab samples. Fecal samples had significantly higher relative abundance of (A) *Peptostreptococcaceae*, (B) *Akkermansiaceae*, (C) *Bacteroidaceae*, compared to grab samples. Also, there was significantly lower relative abundance of (D) *Veillonellaceae*, (E) Bacteroidales_BS11_gut_group and (F) *Spirochaetaceae* compared to grab samples. Points are the estimated relative abundance and bars are a 95% prediction interval for each cow on different days of sampling.

Based on the DPCoA findings, the phyla Spirochaetes and Actinobacteria also played an important role in distinguishing feces from grab samples (Fig 1A and 4B). In the phylum Spirochaetes, there were 10 ASVs, all of which were from the genera Treponema_2, that had significantly lower relative abundance in fecal samples compared with grab samples. Within the phylum Actinobacteria, there were 4 ASVs in the genera *Olsenella*, 5 ASVs in *Atopobium*, 7 ASVs in the genera *DNF00809*, and 1 ASV assigned to *Raoultibacter*, which were all significantly lower in relative abundance compared with grab samples.

### Specific community differences between grab and stomach tube samples

Oral stomach tube samples were composed of 20 phyla, 65 orders, 98 families, and 236 genera. There were 255 ASVs found in grab samples that were not found in the stomach tube samples. Likewise, 404 ASVs in stomach tube samples were not present in the grab samples. There were 3,615 ASVs that were in common between stomach tube and grab samples. Three families *Rhodobacteraceae*, *Bacteriovoracaeae*, and *Spirosomaceae* were found in grab samples, but were not present in stomach tube samples. The 5 families found in stomach tube samples, but not in grab samples were *Cellvibrionaceae*, *Neisseriaceae*, *Bifidobacteriaceae*, *Micrococcaceae*, and *Solirubrobaceraceae*.

In addition to the taxa not found in a particular sample type, there were 13 families, 43 genera, and 199 ASVs significant differentially abundant between stomach tube and grab samples. *Lachnospiraceae*, *Ruminococcaceae*, *Prevotellaceae*, and *Erysipelotrichaceae* were the most common families to have significant differentially abundant ASVs in stomach tube versus grab samples. The relative abundance of 39 ASVs in the family *Lachnospiraceae* were significantly lower while 15 were significantly higher in comparison to grab samples. At the genus level, 15 genera in the family *Lachnospiraceae* were significantly lower in abundance, while *Blautia*, *Acetitomaculum* and *Howardella* were the only genera that had higher relative abundance (S2 Fig). While *Ruminococcaceae* in stomach tube samples was not significantly different from grab samples at the family level (Fig 5B), eight genera in this family were significantly higher in relative abundance between the two sample types. *Prevotellaceae* in stomach tube samples was not significantly different from grab samples at the family level (*P* = *0.055*; Fig 5A), but at the genus level, two were significantly lower and one significantly higher. Three genera in the family *Erysipelotrichaceae, Catenisphaera*, *Erysipelotrichaceae* _UCG-009, and *Erysipelotrichaceae* _UCG-004 were all significantly higher in stomach tube compared with grab samples. The only assigned genera in the family *Fibrobacteraceae*, *Fibrobacter*, was significantly lower in abundance in stomach tubes compared to grab samples (S1 Fig). The genus *Streptococcus* had significantly higher relative abundance compared with grab samples (Fig 6).

The only genus in the phylum Euryarchaeota that had significant differences in abundance in samples from the stomach tube as compared with those collected from the rumen was *Methanobrevibacter*. This genus was significantly higher in stomach tube samples. At a finer resolution, there were only four ASVs assigned to *Methanobrevibacter* and one ASV assigned to *Methnomethylophilaceae* that were significantly higher in abundance in stomach tube samples compared with grab samples. However, at the family level three methanogenic families, *Methanomethylophilaceae*, *Methanobacteriaceae* and *Methanocorpusculaceae*, were not significantly different between the two sample types.

### Comparing sub-fractions of the grab sample

Grab samples of rumen contents were placed in cheesecloth and squeezed to create the liquid strained sample and the solid particulate sample. There were 283 ASVs found in the grab sample that were not identified in the liquid strained samples. Conversely, there were 3,587 ASVs found in common between grab samples and liquid strained samples.

Based on the DPCoA, separation of liquid samples from other rumen sample types was driven in part by taxa from the phylum Kiritimatiellaeota (Fig 4). ASVs in this phylum were only assigned down to the order level with all ASVs assigned to WCHB1-41. Seventeen ASVs from Kiritimatiellaeota were significantly higher in liquid samples compared with grab samples while these ASVs were not significantly different in solid samples versus grab samples.

In addition to Kiritimatiellaeota, the DPCoA suggested that the families *Lachnospiraceae* and *Prevotellaceae* were also a major cause of differences between liquid and grab samples (Fig 4). Differential abundance testing found that indeed *Lachnospiraceae*, *Ruminococcaceae*, and *Prevotellaceae* were the most common families to have significant differentially abundant genera in liquid strained versus grab samples. *Lachnospiraceae* was significantly lower in liquid samples compared with grab samples (*P* < 2.0×10^−16^; Fig 5C). Liquid samples had the most significant differently abundant genera with 22 that had lower relative abundance compared with grab samples and three with higher. One of these genera with significantly higher relative abundance was *Howardella*, which was also higher in relative abundance in the stomach tube samples. Liquid unstrained and liquid strained samples had significantly higher relative abundance in *Prevotellaceae* than grab samples (Fig 5A). Within that family there was higher relative abundance of the genera *Prevotella_1*, *Prevotellaceae_UCG-003, Prevotellaceae_UCG-001* and lower relative abundance of *Prevotellaceae_NK3B31_group* (*P* ≤ 0.01; S2 Fig). In the family *Ruminococcaceae*, there were 7 genera with significantly lower relative abundance.

Liquid samples were also differentiated from grab samples by a significantly lower abundance of Actinobacteria, specifically the family *Eggerthellaceae*, and significantly higher abundance of Lentisphaerae and Cyanobacteria (Fig 1C). ASVs in the phylum Cyanobacteria were all within the order *Gastranerophilales* and were not classified any lower. Likewise, ASVs in the phylum Lentisphaerae were only assigned to the family *Victivallaceae* which were significantly lower in abundance in liquid samples compared with grab samples (Fig 6).

In liquid strained samples there were roughly an equal number of ASVs assigned to the genera *Methanobrevibacter* that were significantly higher and lower in relative abundance compared with grab samples (Fig 6). Therefore at the genus level there was not a significant difference observed in the abundance of the genera *Methanobrevibacter*. Also, in the same phylum Euryarchaeota, there was significantly lower relative abundance of *Methanosphaera* in liquid strained samples when compared with grab samples.

## Discussion

While other studies looked at differences in the rumen microbiome due to rumen sampling method, they usually involved different diets and did not include all the sampling methods presented in the current study. As diet is an important factor that affects the rumen microbiome, we choose to keep the diet consistent during the study to fully investigate the differences between sampling methods. To the authors’ knowledge, this is the first study to compare rumen sampling methods utilizing ASVs rather than OTUs. Therefore, this study has the advantage of identifying AVS that are comparable across studies, which will improve the reproducibility of sequencing studies of the rumen [44].

Kim et al. detected 19 bacterial phyla in the rumen with Firmicutes (57.8%), Bacteroidetes (26.7%) and Proteobacteria (6.9%) in greatest abundance with the remainder of the 16 phyla less than 3% of the total sequences [45]. In the present study, 21 phyla were identified in the grab sample and only three phyla were over 3% abundance: Firmicutes (64.3%), Bacteroidetes (20%) and Spirochaetes (4.1%). This differed from fecal samples where the top four phyla were Firmicutes (61.2%), Bacteroidetes (32.1%), Verrucomicrobia (1.3%) and Proteobacteria (1.1%) with the remainder of the phyla observed at less than 1% mean relative abundance (Fig 1A and 1B). The relative abundance of Firmicutes and Bacteroides in the fecal samples were similar to what Wong et al. found in fresh manure, but they found Actinobacteria among the top four phyla rather than Verrucomicrobia [46].

The day a sample was collected did not affect the number of species sampled and did not impact the abundance of *Prevotellaceae, Ruminococcaceae* and *Lachnospiraceae.* These observations agree with previous work that found there was little day-to-day variation in both the solid and liquid fraction of rumen samples from the same animal [47]. Previous work found differences between breeds [6,7], while others found minimal to no influence of breed [8,48]. As we only had one Jersey as part of this study we are unable to determine the impact of breed on the community composition.

### Diversity

In the present study, fecal samples had lower richness when compared to grab samples. This is in agreement with a study that used Faith’s Phylogenetic Diversity to compare samples from esophogeal tubing or feces of beef calves [49]. The same result was found using the number of ASVs present in fecal compared to rumen condense after slaughter [50]. Similar to fecal samples, we found that samples collected via the esophageal tube had lower richness than grab samples. Such a finding was expected as microbes adhered to particles would be in low proportion or excluded in the stomach tube sample, even though the tube used did not have a screen. Using a stomach tube without a screen allowed the collection of small size particulates only, whereas the grab samples included small to large particulate sizes. Our finding contradicts Paz et al. who reported no difference in richness between a rumen sample collected from a rumen cannula compared with a sample collected via esophageal tube [8]. However, in Paz et al., solid particles that adhered to the metal strainer of the esophageal tube were recovered and added to the esophageal sample to create a sample that was “more adequately representative of the rumen content”, which suggests the authors acknowledge that a sample collected by an esophageal tube that did not contain particles would not represent rumen contents. However, the research did not address this suggestion by analyzing the rumen contents collected with a stomach tube without the added solids.

Our work also differed from that of Ji et al. who reported the diversity of the bacterial population was not affected by sample type [51]. Samples in their study included rumen digesta collected from a cannula that was squeezed through cheese cloth to create a liquid and a solid fraction for comparison with rumen digesta from a cannula. However, we determined that both liquid sample types did not have significant differences in the number of taxa observed compared to grab samples, while solid samples had significantly lower estimated species than grab samples. The work of Weimer et al. (2017) used a sample cup to collect 100 ml of digesta from the medio-ventral region of the rumen followed by squeezing through cheese cloth to create a liquid and a solid sample [31]. While this study found that community diversity and community richness were greater in solids than liquid, our data showed the opposite. Greater richness in liquid samples could potentially be explained by the greater relative abundance of *Prevotellaceae*, the most abundant species in the rumen, compared with the estimated number of species in solid samples. Jewel et al. found liquid samples to have higher richness than solid samples in agreement with our data [52].

Some of these discrepancies are in part due to differences in the metric used to estimate richness. All these previous studies reported Chao1 as a measure of richness, but the current study used breakaway to estimate richness. Many alpha diversity estimates that are ubiquitous in the literature are highly biased and require statistical adjustments to address this bias, which Chao1 does not [40,41]. Further, the strong negative bias of Chao1 is even further increased by the use of rarefying as a means of normalization in the previous studies [53]. It is true that Dr. Anne Chao proposed that Chao1 could be a useful metric for datasets that skewed toward low-abundance classes as microbiome data does; however, these low abundance counts aren’t reliable due to sequencing platform and PCR errors. Breakaway addresses some short comings of Chao1 by providing an estimate of variance of richness estimates for hypothesis testing, estimating the number of missing taxa, and adjusting the richness estimate accordingly (bias correction) to provide a more accurate estimate of richness [41]. While this approach produces large error bars, the breakaway estimate provides a more accurate reflection of the uncertainly associated with estimating a true value that can never be known (Fig 2B).

### Bacterial populations

#### Rumen samples

Based on the exploratory analysis with the DPCoA, differences between rumen liquid strained samples and other rumen samples types were driven mainly by *Lachnospiraceae*, *Prevotellaceae* and Kiritimatiellaeota. *Lachnospiraceae* was significantly lower in liquid samples and *Prevotellaceae* had significantly higher relative abundance compared with grab samples (Fig 5A and 5C). Other studies that examine differences between the microbial communities in liquid and solid phases have reported both *Lachnospiraceae* and *Ruminococcaceae* in higher and in lower abundance in the liquid samples compared with the solid [19,54]. These conflicting results could be due to the different diets used in these studies. Animals on all forage diets had higher abundances of both families in liquid phase, while cattle on a diet with a forage to concentrate ratio of 70:30 had lower abundances of these families in the solid phase [19,54]. Lower resolution of the taxa might lend clues as to the cause of these differing results.

In agreement with our study, others have found that *Prevotellaceae* were most abundant in liquid phase compared with solid phase and the dominant family in the liquid fraction [19,21,22]. *Prevotella sp.* are capable of degrading a wide variety of substrates including pectin, hemicellulose, protein, fatty acids, and starch [55]. Readily fermentable carbohydrates including sugars and soluble fiber in the liquid fraction likely support the presence of *Prevotella*. Thus, the lower abundance of *Prevotella* in samples with increased solid fraction, including grab samples and solid strained was logical.

Our data show that ASVs from Kiritimatiellaeota had significantly higher abundances in liquid strained samples, but these ASVs were not significantly differentially abundant in solid versus grab samples (Fig 1A and 1C). These data are in agreement with a study that found Kiritimatiellaeota in higher proportion in the liquid compared with the solid phase of a yak rumen [56]. Additionally, an order in this phyla, WCHB1-41, was identified to be part of the “core microbiome” in liquid samples from the rumen [57]. Kiritimatiellaeota was found in rumen samples and was in higher abundance from samples of higher methane producers making it a potentially important microbe to understand in order to possibly reduce methane emissions [58]. Bioinformatic analysis has hypothesized that this phyla uses sodium for a coupling ion to generate the electrochemical gradient to produce ATP, rather than the typical H^+^ [59]. Therefore, in circumstances when concentrations of H^+^ are relatively lower, as when methane emission are high, this phylum could have a competitive advantage of using sodium as a coupling ion. The role of this rumen microbe has yet to be understood and our data demonstrates that for investigators interested in elucidating the role of this microbe in the rumen ecosystem, samples can be enriched with Kiritimatiellaeota by filtering rumen samples through cheese cloth.

#### Stomach tube samples

In a previous study, when sampling was done by either rumen cannula or esophageal tube *Prevotellaceae*, *Lachnospiraceae* and *Ruminococcaceae* were the predominate families regardless of the sampling method [8]. Importantly, these authors made a point to include particles attached to the strainer to capture a representative sample in the rumen. Similarly, in the present study *Prevotellaceae* and *Ruminococcaceae* (Fig 5A and 5B) were not significantly different at the family level, while *Lachnospiraceae* was significantly lower in stomach tube samples (Fig 5C). The lower relative abundance of *Lachnospiraceae*, specifically the genera *Butyrivibrio* and *Coprococcus*, in samples collected by esophageal tube rather than through a rumen fistula was also determined in another study (S2 Fig) [60]. However, at a finer resolution our data showed that these three families had the most significant differentially abundant ASVs when comparing the stomach tube and grab samples.

In agreement with De Menezes et al. who found *Fibrobacter* and *Spirochaetes* in the solid fraction, the only assigned genera in the family *Fibrobacteraceae*, *Fibrobacter*, was significantly lower in abundance in stomach tube samples compared with grab samples (Fig 1C, 6 and S2 Fig) as was the family *Spirochaetaceae* (Fig 1C and 6) due to a lower abundance of the genus *Treponema* (S2 Fig) [61]. Initially, we hypothesized the lower abundance of *Fibrobacter* species in stomach tube samples would largely be driven by the exclusion of fibrous particles in the sample as *Fibrobacter* facilitates cellulose degradation in the rumen [62–64]. However, significantly lower abundances of the family *Fibrobacteraceae* and *Fibrobacter* at the genus level were seen in solid and liquid unstrained samples compared to grab samples (Fig 6 and S2 Fig). Alternatively, the differences could be attributed to location of rumen sampling.

Another fiber adherent bacterium *Ruminococcus flavefaciens* (contained in genus Ruminococcus_1) did follow the expected pattern of significantly lower abundance in stomach tube and liquid samples and significantly higher abundance in solid samples compared with grab samples (S2 Fig). The different distribution of these to cellulolytic species could be reflective of their differential preferences for particular plant tissues, for example structural polysaccharides of the cell wall, as a growth substrate [65]. For studies that are interested in fibrolytic bacteria such as *Fibrobacter*, straining the liquid out of the sample does not enrich for these bacteria, but rather seems to disrupt these communities. Therefore, our data suggests that grab samples are the best option for examining these populations.

An important phylum in defining stomach tube samples was Fusobacteria, which was significantly higher in abundance in stomach tube samples compared with grab samples (Fig 1C). This difference was driven by the genus *Fusobacterium* (S2 Fig) and to the authors’ knowledge this difference between stomach tube and rumen sampling methods has not been previously reported. *Fusobacterium necrophorum* is an important target species for improving rumen efficiency as it degrades lysine, whose dietary deficiency is the most likely to limit milk production [66,67]. In addition, *F*. *necrophorum* was reported to be an opportunistic pathogen that causes liver abscesses in feedlot cattle [68,69]. Our data have identified a previously unreported difference between rumen and stomach tube samples that would enable monitoring of this important genus with stomach tube sampling and has implications for both dairy and beef cattle.

Stomach tube samples more closely reflected liquid samples, but stomach tube samples were highly variable (Fig 3 and S1 Fig). This high variability in microbial community could reflect the fact that the stomach tube did not have a screen, therefore the solid contribution to the stomach tube sample was also highly variable. There were 3,615 ASVs that were in common between stomach tube and grab samples. Two families, *Rhodobacteraceae* and *Spirosomaceae* were found in grab, liquid strained and liquid unstrained samples, but were not present in stomach tube samples. However, *Solirubrobacteraceae* was found only in stomach tube samples. These differences could reflect differences in the location of the tube placement (cranial ventral) compared with the sampling the rumen from the cannula (central rumen).

Taken together, these data suggest that stomach tube samples could be reflective of rumen samples provided some solid particulate are included and attempts are made to place the tube at a consistent depth. Despite following these precautions, researchers should expect these samples to be more variable than grab samples and increase their sample size accordingly.

#### Feces vs rumen

In the current study, as anticipated, fecal samples were not representative of the microbial community of the rumen. The differences between fecal and rumen samples were driven by differences in two Firmicute families: *Ruminococcaceae* and *Lachnospiraceae* (Fig 4). Indeed, it was found that there was significantly higher abundance of *Ruminococcaceae* (Fig 5B) and significantly lower abundance of *Lachnospiraceae* in feces (Fig 5C). Similarly, Noel et al. found the abundance of *Ruminococcaceae* to be much higher in feces compared with rumen samples [70]. However, they found no difference in the abundance of *Lachnospiraceae*. A recent preprint found strikingly similar relative abundances of top three most abundant families in feces from dairy cattle: *Ruminococcaceae* (34.9% compared to our 40.7%), and *Rikenellaceae* (11.6% compared to our 15.7%) and *Lachnospiraceae* (6.8% compared to our 7.7%) [71]. These data show that *Ruminococcaceae* is typically found in higher abundance in feces, while fecal *Lachnospiraceae* will have lower abundance than the rumen population.

Both *Lachnospiraceae* and *Ruminococcaceae* are also members of the human gastrointestinal tract and have multiple glycoside hydrolases (GH) and carbohydrate-binding modules (CBM) that allow utilization of complex plant material, and transport degradation products of various sizes and compositions [72]. Their differences in abundance between the rumen and fecal samples was likely a reflection of their specialization in degrading the various types of substrates present in these two niches. As both families contain butyrate producers, the shift in these families could represent a change in the major sources of butyrate in the rumen compared with the lower colon. The reader should note that there are discrepancies in the literature as to the taxonomy of genera in *Lachnospiraceae* [73]. Of note is a prominent butyrate producer *Eubacterium rectale* that is cited as belonging to both *Eubacteriaceae* and *Lachnospiraceae*, despite its placement on a 16S rRNA gene tree near recognized members of *Lachnospiraceae* [74]. These inconsistencies can make appropriate comparisons at the level of family across studies difficult.

In addition, to the families that drove the major differences between rumen and feces, other families were also found to be differentially abundant between these two sample types. There was significantly higher abundance of *Akkermansiaceae* in feces compared with grab samples (Fig 6B). Until 2016, *Akkermansiaceae* only contained the species *Akkermansia muciniphila*, when a novel strain, *Akkermansia glycaniphila*, was isolated from the feces of a reticulated python [75]. Muciniphila means “mucin-loving” in Latin and as its name suggests *A. muciniphila* is a mucin-degrader, which produces acetate and propionate from mucin fermentation [76]. This species is known to be one of the most abundant in the human colon making up 0.5-5% of the total bacteria, which was in agreement with the relative abundance we observed (Fig 7B) [77,78]. Other studies have also noted the higher abundances of *Akkermansia* in feces compared with rumen samples [28,79]. In humans, *A. muciniphilia* had a protective effect against obesity and played a role in both glucose and lipid metabolism [80,81]. *Akkermansia* also had anti-inflammatory effects that were in part mediated through a membrane specific protein that interacted with the toll-like receptor-2 and improved gut-barrier function when given orally [82]. Due to the role of *A. muciniphilia* in regulating intestinal inflammation and fat deposition, a better understanding of its function in cattle could identify methods to improve weight gain in cattle.

Taken together, fecal samples are not an accurate representation of rumen samples as they have differences in the abundance of predominant families in the phyla Firmicutes and Bacteroidetes. Fecal samples differed from those taken from the rumen as they had significantly lower relative abundance of *Lachnospiraceae, Christensenellaceae, Prevotellaceae, Fibrobacter* and *Treponema* (Fig 5A and C, 6, 7 and S2 Fig). Also, fecal samples had significantly higher relative abundance of *Ruminococcaceae, Rikenellaceae* and *Akkermansia* compared with grab samples (Fig 5B, 6, 7 and S2 Fig). Researchers can access the freely accessible data found at https://doi.org/10.5281/zenodo.4026849 to determine how sampling methods might affect the abundance of their microbe of interest.

### Archaeal populations

#### Feces vs rumen

Methanogens are an important functional group within the rumen as their use of H_2_ to reduce CO_2_ to methane (CH_4_) removes H_2_ from the rumen that is generated during fermentation of carbohydrates [83,84]. Methane has a global warming potential 28-34 fold higher than CO_2_ over 100 years, and therefore its mitigation is important to reducing the environmental impact of animal agriculture. Additionally, methane production is energy inefficient, resulting in a 2-12% loss in gross energy intake in cattle [85]. There is very limited data on differences between the archaeal populations in the rumen compared with the feces, as a majority of studies solely focus on the rumen population.

One study that has examined both the rumen and fecal populations of archaea of Nelore cattle was conducted by Andrade et al. [50]. Like this present study, Andrade et al. also utilized DADA2 to identify ASVs and assigned taxa with the SILVA database v132; however, they used different primers that are specific for archaea and bacteria rather than universal primers and classified archaeal sequences using the Rumen and Intestinal Methanogen database (RIM-DB). Together these choices allowed Andrade et al. to classify archaeal ASVs down to the species level, which contrasted with this present study where methanogenic ASVs were only classified down to the genus level. Other than *Methanobrevibacter* and *Methanosphaera*, the other archaeal genera that this present study and Andrade et al. identified were different. Our data contained *Methanocorpusculum*, *Methanimicrococcus* and Candidatus *Methanomethylophilus* while Andrade et al. observed *Methanomicrobium*. Both studies found that *Methanobrevibacter* and *Methanosphaera* were found in both the rumen and feces; however, there were differences in the relative abundances of the main genera. In contrast to Andrade et al. we found significantly lower relative abundance of *Methanobrevibacter* in fecal samples compared with samples from the rumen. Despite using similar methods there is not clear agreement as to the differences in abundance of genera and which genera are present in the two populations.

As an alternative to 16S rRNA gene sequencing, the mcrA gene can be sequenced to study methanogens [86,87]. The mcrA gene encodes the α-subunit of the methyl coenzyme M reductase, which catalyzes the last step of methanogenesis and is conserved among all methanogens [88]. A study that used mcrA amplicon sequencing found that the most abundant genera in manure was *Methanocorpusculum* while in the rumen it was *Methanobrevibacter* [89]. While we found *Methanocorpusculum* in our fecal samples it was a minor genus and the discrepancy is most likely explained by differences in the gene amplicon sequences. Taken together these data suggest *Methanobrevibacter* is a dominant archaeal genus in both the rumen and fecal populations. The lack of data comparing the rumen and fecal populations suggest that further research is required to understand the archaeal populations.

#### Rumen samples

In the present study relative abundance of archaeal families was similar across rumen samples, both liquid and solid phases, with wide variation in the relative abundance of *Methanocorpusculaceae* (Fig 6). In contrast, Bowen et al. found methanogens to be more abundant in the solid phase [19]. Our data more closely agree with de Mulder et al. who found similar abundance in samples including solid, rumen liquid, and liquid [54]. When we examined the archaeal ASVs in our data at the genus level, *Methanosphera* was significantly lower in relative abundance in liquid samples compared with grab samples. This is in agreement with previous studies that found *Methanosphera* was more abundant in the solid phase, rather than the liquid phase [19,54]. As a whole these data suggest that the collective abundance of methanogens was similar between solid and liquid phases, but that *Methanosphera* are found at higher abundance in the rumen liquid. Studies evaluating feed additives or diet alterations to modulate methanogen populations in the rumen should consider including the liquid fraction of rumen fluid to capture changes in the abundance of *Methanosphera*.

At the family level three methanogenic families, *Methanomethylophilaceae*, *Methanobacteriaceae* and *Methanocorpusculaceae*, were not significantly different between the grab sample and samples acquired via a stomach tube (Fig 6). However, there were 4 ASVs assigned to *Methanobrevibacter* that were found to be significantly higher in abundance in stomach tube samples. This is a paradoxical finding as stomach tube samples typically have more liquid than solid particles in them and we previously noted that *Methanosphaera* was in higher abundance in liquid samples. As the coefficient for the difference in relative abundance of *Methanobrevibacter* is low (0.1-0.5), we believe that in practice with higher numbers of animals this difference would be negligible.

Many of the differences described thus far have focused on the major genera *Methanobrevibacter* and *Methanosphaera*, which are hydrogenotrophic methanogens. While the hydrogenotrophic pathway for methane production is the most common there are two alternative pathways: methylotrophic and acetoclastic that utilize methylated compounds and acetate, respectively. Thus far, only taxa within the order *Methanosarcinales* have been identified to be capable of acetoclastic methanogenesis [90,91]. An acetoclastic methanogen in our data, *Methanimicrococcus*, was only present in two liquid samples. There was not a strong pattern as to the phase in which this minor genus may be found, and as deep sequencing would be required to determine shifts in its abundance, targeted qRT-PCR would be a better choice to study abundance of this microbe. In addition, there was one ASV assigned to *Methnomethylophilaceae*, a methylotrophic archaeon, that was significantly higher in abundance in stomach tube samples compared with grab samples, although at higher taxonomic levels no differences were found for the family *Methnomethylophilaceae*.

Taken together these results demonstrate that stomach tubing would likely provide a representative community of major populations of methanogens, *Methanosphaera* and *Methanobrevibacter*, compared with grab samples. For minor populations accurate surveys would require more targeted techniques, such as qRT-PCR or mcrA sequencing. While this study added to an understanding of how sampling methods will potentially impact archaea populations observed, it should not be considered a comprehensive evaluation of the microbial communities. Specific archaeal primers and qRT-PCR could be used to clarify discrepancies between this study and past work. However, for those evaluating archaeal communities with 16S rRNA gene sequencing, this study can serve as a guide to help in study design to improve the chances of capturing an accurate picture of the taxa of interest.

## Acknowledgements

We thank the UC Davis DNA Technologies Core for sequencing services. We appreciate the work of the UC Davis Dairy manager Douglas Gisi and Carlyn Peterson that helped during sample collection.

## Author contribution statement

JH analyzed, interpreted data, and wrote manuscript. ML collected samples, prepared libraries, aided in experimental design and edited manuscript. ED designed the experiment, assisted in sample collection helped prepared manuscript. EAM edited manuscript.

## Supporting information

**S1 Fig. Double principal coordiant analysis of the Bray-Curtis distance after removal of the phyla Bacteroidetes and Firmicutes from the dataset.** DPCoA is a phylogenetic ordination method and that provides a biplot representation of both (A) samples and (B) taxonomic categories. The 1^st^ axis separtes liquid strained samples from other rumen sample types while the 2^nd^ axis discrimates fecal from rumen samples. Samples that have larger scores on the 1^st^ axis have more taxa from the phylum Kiritimatiellaeota and less taxa from the phylum Euryarchaeota. Likewise, samples with higher scores on the 2^nd^ axis have more taxa from the family Akkermansiaceae and less taxa from the families Fibrobacteraceae and Spirochaetaceae. To faithfully reflect the variance in the coordinates, the height-to-width ratio was based on the ratio between the corresponding eigenvalues.

**S2 Fig. Significant genera that are differentially abundant across sample type graph as coefficients with a 95% confidence interval calculated from the corncob model.** Taxa with negative coefficients for a sample type are expected to have a lower relative abundance when compared to the grab samples while positive coefficients suggest a higher relative abundance in that sample type compared to grab samples. Taxa are presented with phylum, family, genus and species to the lowest assigned level.

